# CAR Tregs mediate linked suppression and infectious tolerance in islet transplantation

**DOI:** 10.1101/2024.04.06.588414

**Authors:** Christine M. Wardell, Vivian C.W. Fung, Eleanor Chen, Manjurul Haque, Jana Gillies, Justin A. Spanier, Majid Mojibian, Brian T. Fife, Megan K. Levings

## Abstract

Regulatory T cells (Tregs) have potential as a cell-based therapy to prevent or treat transplant rejection and autoimmunity. Using an HLA-A2-specific chimeric antigen receptor (A2-CAR), we previously showed that adoptive transfer of A2-CAR Tregs limited anti-HLA-A2 alloimmunity. However, it was unknown if A2-CAR Tregs could also limit immunity to autoantigens. Using a model of HLA-A2^+^ islet transplantation into immunodeficient non-obese diabetic mice, we investigated if A2-CAR Tregs could control diabetes induced by islet-autoreactive (BDC2.5) T cells. In mice transplanted with HLA-A2^+^ islets, A2-CAR Tregs reduced BDC2.5 T cell engraftment, proliferation and cytokine production, and protected mice from diabetes. Tolerance to islets was systemic, including protection of the HLA-A2^negative^ endogenous pancreas. In tolerant mice, a significant proportion of BDC2.5 T cells gained FOXP3 expression suggesting that long-term tolerance is maintained by *de novo* Treg generation. Thus, A2-CAR Tregs mediate linked suppression and infectious tolerance and have potential therapeutic use to simultaneously control both allo- and autoimmunity in islet transplantation.

**One Sentence Summary:** Alloreactive chimeric antigen receptor-engineered regulatory T cells limit diabetogenic T cell engraftment and function to prevent type 1 diabetes.

## INTRODUCTION

FOXP3^+^ regulatory T cells (Tregs) suppress dysregulated inflammation and maintain immune homeostasis. Given their tolerogenic properties, there is significant interest in using Tregs as a cell-based therapy in applications such as transplantation and autoimmunity to reduce reliance on pharmacologic immunosuppression. Clinical trials of Treg therapy to date have primarily tested autologous polyclonal, *ex vivo*-expanded Tregs showing excellent safety and tolerability (*1, 2*). However optimal clinical efficacy is thought to depend on antigen specificity, with multiple pre-clinical studies showing that Tregs enriched or engineered for disease-relevant antigen specificity are more therapeutic than their polyclonal counterparts (*3–5*).

Approaches to enrich or engineer Tregs for antigen specificity include stimulation with relevant antigens during expansion (*6–8*), or introduction of antigen-specific T cell receptors (TCRs) or chimeric antigen receptors (CARs) (*3, 4, 5*). Focusing on CARs, we and others developed CARs which recognize HLA-A2 (A2-CAR), a commonly mismatched protein in organ transplantation. Tregs engineered to express an A2-CAR suppress anti-HLA-A2-directed alloimmunity significantly better than polyclonal Tregs in multiple humanized and mouse transplant models (*9–16*). This pre-clinical work led to rapid clinical translation and two ongoing clinical trials of A2-CAR Tregs in kidney and liver transplantation ((*17*) NCT04817774 for kidney, NCT05234190 for liver).

A fundamental premise of Treg therapy is its potential to induce long-lasting tolerance via bystander or linked suppression, as well as infectious tolerance, concepts introduced by Dr. H. Waldmann through studies in transplantation. The concept of linked suppression describes situations where tolerance towards antigen “A”, is transferred to antigen “B” when co-expressed with antigen “A” (e.g. A x B). Tolerance to Antigen “B” subsequently becomes independent of antigen “A” co-expression (*18*). More recently, a less-defined version of this concept has come to be known as bystander suppression, which generally refers to simultaneous suppression of immunity to two co-expressed antigens, but stops short of demonstrating independent tolerance to the bystander antigen. Infectious tolerance refers to a Treg-mediated, long-lived increase in the ratio of Tregs to effector T cells (Teff) (*19*). Although the exact mechanisms of infectious tolerance remain unclear, the process likely depends on a combination of Treg functions including contact-dependent Teff and antigen presenting cell (APC) inhibition, TGF-β-driven FOXP3 expression and environmental metabolic disruption (*18, 20–25*).

The potential for therapeutic Tregs to control immunity to multiple antigens and have a long-lived tolerogenic effect via expansion of endogenous Tregs is highly desirable in the cell therapy context. Through these mechanisms, a single, potentially transient, dose of Tregs could have a long-lasting effect. However, evidence supporting the ability of CAR-engineered Tregs to induce linked/bystander suppression and infectious tolerance was lacking. Here we tested the hypothesis that A2-CAR Tregs could induce long-lasting tolerance to antigens distinct from HLA-A2 using a mouse model of islet transplantation.

## RESULTS

### Generation of A2-CAR Tregs from non obese diabetic mice

Islet transplantation can provide a functional cure for type 1 diabetes (T1D) (*26*), but success is limited by rejection mediated by both alloreactive and re-current autoreactive T cells (*27*). To test if A2-CAR Tregs mediate linked/bystander suppression to control autoimmunity, we established a model of HLA-A2^+^ islet transplantation in a diabetogenic T cell adoptive transfer model of T1D.

We first generated polyclonal and antigen-specific A2-CAR Tregs from non obese diabetic (NOD) mice by sorting and expanding CD4^+^GFP^+^ cells from the spleen and lymph nodes of NOD.FOXP3-GFP (Thy1.2^+^) mice (*28*). Three days after activation, Tregs were transduced with a retrovirus encoding either the transduction marker Thy1.1 alone (referred to as polyclonal Tregs hereafter), or both the A2-CAR and Thy1.1 (referred to as A2-CAR Tregs hereafter) (*9*) (Fig 1A). The resulting expanded Tregs expressed high levels of both lineage-defining transcription factors, FOXP3 and HELIOS (mean double positive = 82% for polyclonal and = 86% for A2-CAR; Fig 1B), as well as Thy1.1 (mean = 88% for polyclonal and = 100% for A2-CAR; Fig 1C). For the A2-CAR Tregs, expression of Thy1.1 and the CAR (myc tag) were directly correlated (Fig 1C, left flow plot).

**Fig. 1:**
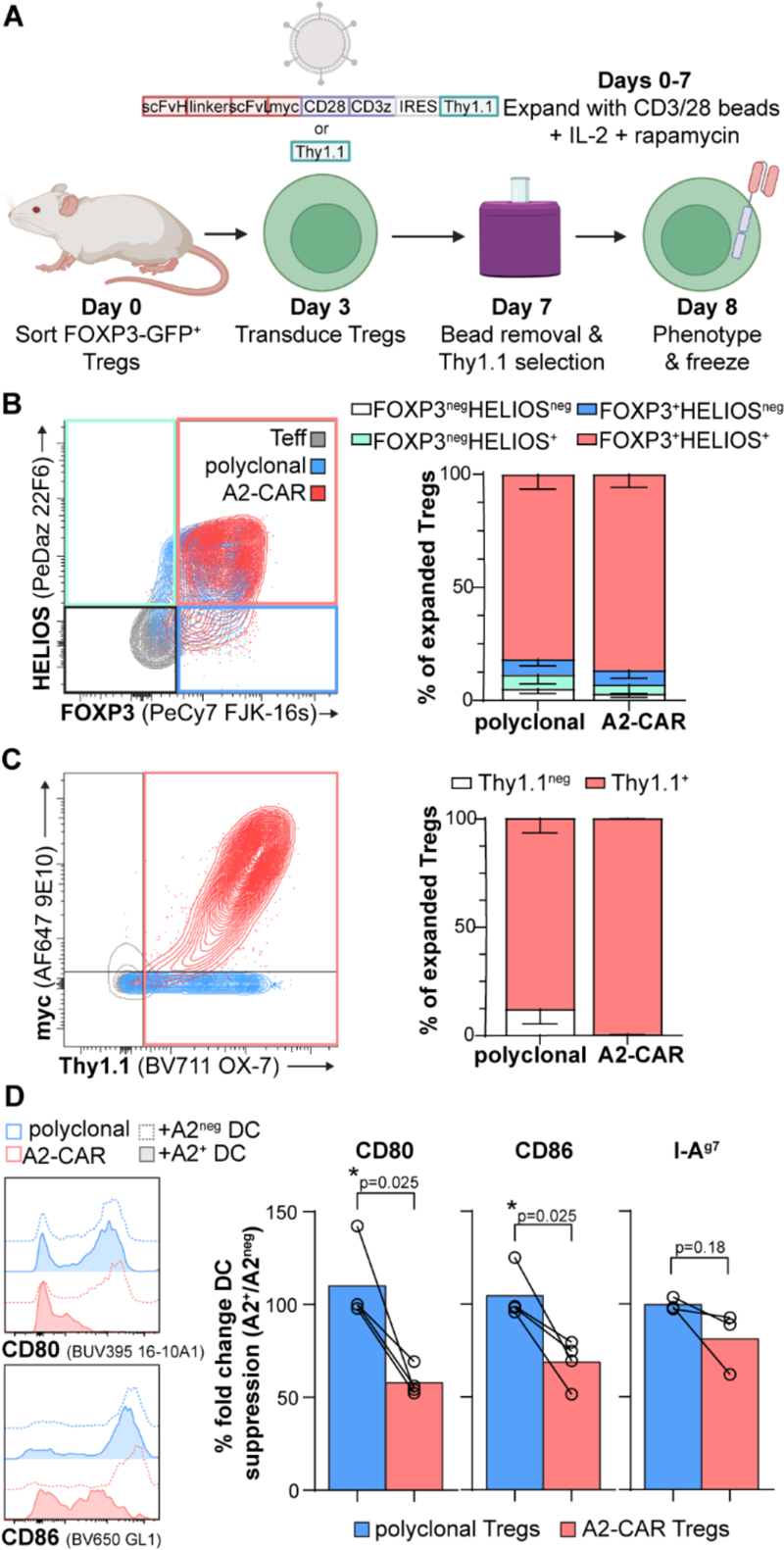
A2-CAR Treg generation from NOD.FOXP3-GFP mice. **(A)** FOXP3-GFP^+^ Tregs were sorted from reporter NOD mice and stimulated in the presence of CD3/CD28 beads, IL-2 and rapamycin. Tregs were retrovirally transduced on day three to express the transduction marker Thy1.1 and the A2-CAR. Beads were removed on day 7, and transduced cells were magnetically selected and rested overnight before being frozen on day eight. Markers of Treg stability, FOXP3 and HELIOS **(B)**, and Thy1.1 expression **(C)**, were quantified on the day of freezing (n=7; mean and SD). **(D)** Expanded Tregs were cultured with HLA-A2^negative^ or HLA-A2^+^ NOD splenic DCs for two days; representative histograms of Treg effect on DC expression of co-stimulatory CD80 (top) and CD86 (bottom) are shown on the left and quantified on the right, along with I-A^g7^ (n=3-4; bar at mean with individual pairs shown; paired t-test).

Linked and bystander suppression are thought to be mediated at least in part through Treg-mediated induction of tolerogenic APCs, via removal of co-stimulatory molecules and MHC molecules (*29*). To test if NOD A2-CAR Tregs influenced APC activity, the former were co-cultured with HLA-A2^negative^ or HLA-A2^+^ NOD splenic dendritic cells (DCs) (*16*). After 48 hours, DCs were analyzed for expression of the co-stimulatory molecules CD80 and CD86, and the NOD MHC class II molecule, I-A^g7^. Whereas polyclonal Tregs had a similar effect on HLA-A2^negative^ and HLA-A2^+^ DCs, A2-CAR Tregs significantly reduced the expression of CD80 and CD86 on HLA-A2^+^ relative to HLA-A2^negative^ DCs, and showed a similar trend for I-A^g7^ (Fig 1D). Thus, NOD A2-CAR Tregs have a heightened, antigen-specific ability to decrease the potential for APCs to activate Teff.

### A2-CAR Tregs activated by HLA-A2^+^ islets protect from autoimmune diabetes

To test if A2-CAR Tregs could suppress autoreactive Teff, we next established a model of islet transplantation and autoimmune diabetes (Fig 2A). 50-150 HLA-A2^+^ NOD islets were hand-picked from pre-diabetic, 6-11 week old mice, transplanted into the left anterior chamber of the eye (ACE) of NSG mice, and allowed to engraft for at least 2 weeks. The ACE is a validated location to study islet-directed auto- and alloimmunity (*30–34*): in NOD mice, ACE islet transplants are subject to autoimmune attack at the same rate as islets in the native pancreas (*30*).

**Fig. 2:**
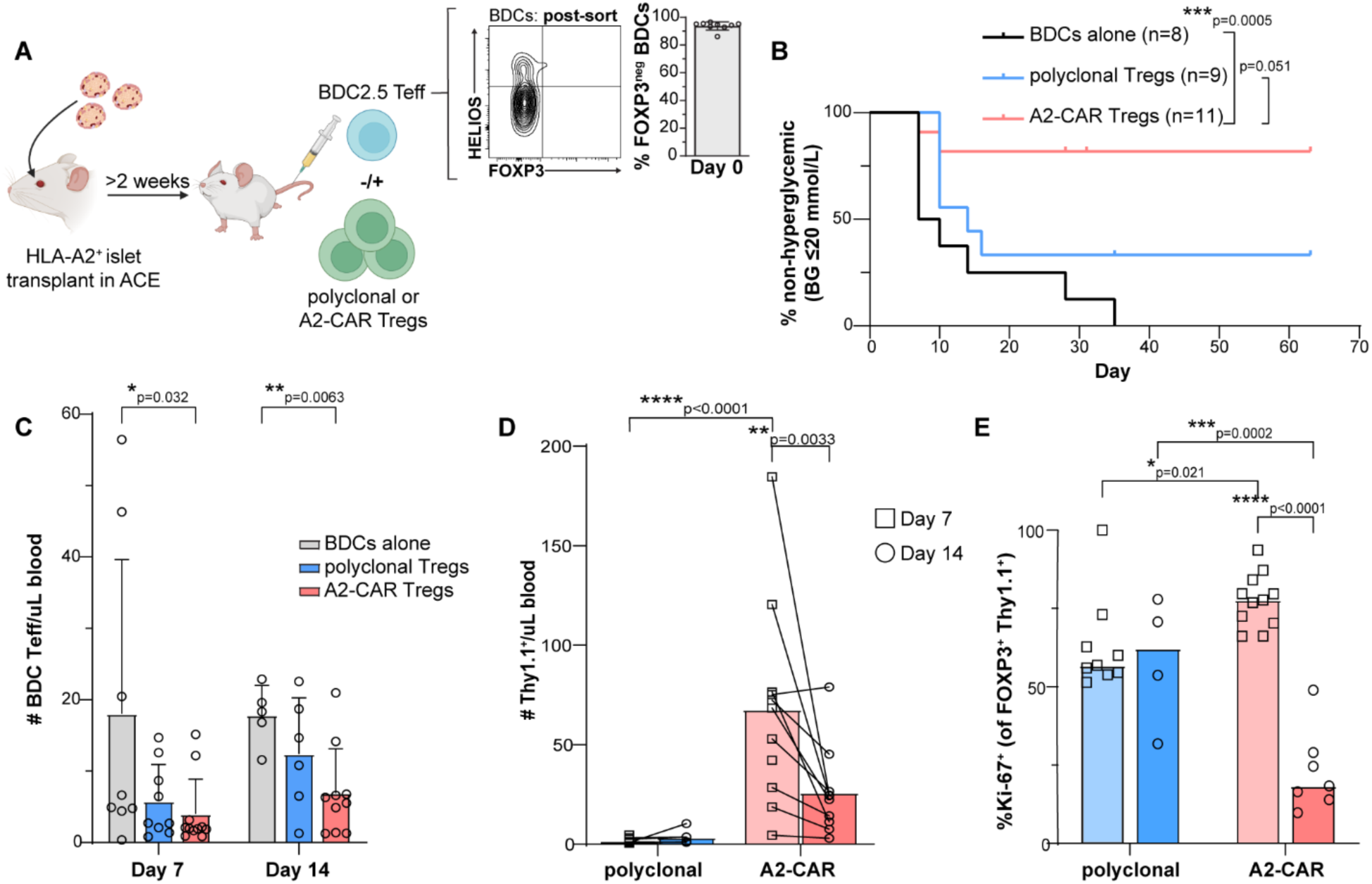
A2-CAR Tregs protect mice with an HLA-A2+ islet transplantation from autoimmune diabetes. **(A)** NSG mice were transplanted with 50-150 islets from HLA-A2^+^ NOD mice in the left anterior chamber of the eye (ACE). At least two weeks post-transplant, mice were injected with FOXP3^negative^ BDC Teffs sorted from NOD.FOXP3-GFP x BDC2.5 mice; representative and average FOXP3 and HELIOS expression is shown (mean ± SD). Some mice were also treated with polyclonal or A2-CAR Tregs (1 BDC2.5 Teff:3 Treg ratio). **(B)** Diabetes-free survival of each treatment group, with ≥20 mM blood glucose reading defining diabetes onset (n=8-11; Mantel-Cox Logrank test; ticks indicate censored mice). **(C)** FOXP3^negative^ BDC2.5 Teff engraftment in blood at days 7 and 14 post-cell injection (n=5-11; mean ± SD; mixed effects analysis with Tukey’s multiple comparisons). **(D)** Absolute number of Thy1.1^+^ Tregs engrafted in blood over time (n=9-11; individual pairs shown with bars at means; mixed effects analysis with uncorrected Fisher’s LSD). **(E)** Frequency of Ki-67-expressing cells within adoptively transferred FOXP3^+^Thy1.1^+^ Tregs in the blood (n=4-11; bar at median; mixed effects analysis with uncorrected Fisher’s LSD).

Diabetogenic BDC2.5 Teff that recognize a hybrid peptide between the insulin C chain and chromogranin A (*35*) were sorted as CD4^+^FOXP3-GFP^negative^ cells from NOD.FOXP3-GFP x BDC2.5 mice to obtain BDC2.5 Teff that were FOXP3^negative^ (Fig 2A). As expected, within a median of 7 days, injection of 50,000 BDC2.5 Teff induced hyperglycemia in HLA-A2^+^ islet recipient mice (Fig 2B). Co-injection with 150,000 polyclonal Tregs protected 33% of recipient mice against hyperglycemia; however, co-injection of the same number of A2-CAR Tregs stably protected most (82%) of the treated mice.

T cell engraftment was tracked weekly in blood using flow cytometry. One- and two-weeks post-T cell injection, A2-CAR Treg-treated mice had a significantly lower number of circulating BDC2.5 Teff compared to untreated mice (Fig 2C). A2-CAR Tregs engrafted in the blood at high levels relative to their polyclonal counterparts (Fig 2D), and experienced a proliferative, presumably antigen-stimulated, burst by day 7 (Fig 2E). These data suggest that expression of the CAR antigen on the transplanted islets activated A2-CAR Tregs to reduce engraftment of the co-expressed diabetogenic peptide-recognizing BDC2.5 Teff population.

### A2-CAR Tregs reduce BDC2.5 T cell effector function

In our model, the endogenous HLA-A2^negative^ pancreas was unmanipulated. This allowed us to evaluate whether A2-CAR Treg treatment reduced the ability of BDC2.5 Teffs to be activated by their cognate antigen even in the absence of co-expressed A2-CAR Treg antigen, which would be suggestive of linked suppression. Seven days post-T cell injection, flow cytometric analysis of the endogenous pancreas revealed that mice which received A2-CAR Tregs had diminished BDC2.5 T cell infiltration in the exocrine pancreas (Fig 3A), and that they were less proliferative (Fig 3B). Moreover, the native pancreas of A2-CAR Treg treated mice had less severe insulitis compared to mice which did not receive Tregs (Fig 3C). This suppressive effect of A2-CAR Treg on BDC2.5 Teffs was systemic as there were also fewer and less proliferative BDC2.5 Teff in the spleens of A2-CAR Treg treated mice (Fig 3D&E).

**Fig. 3:**
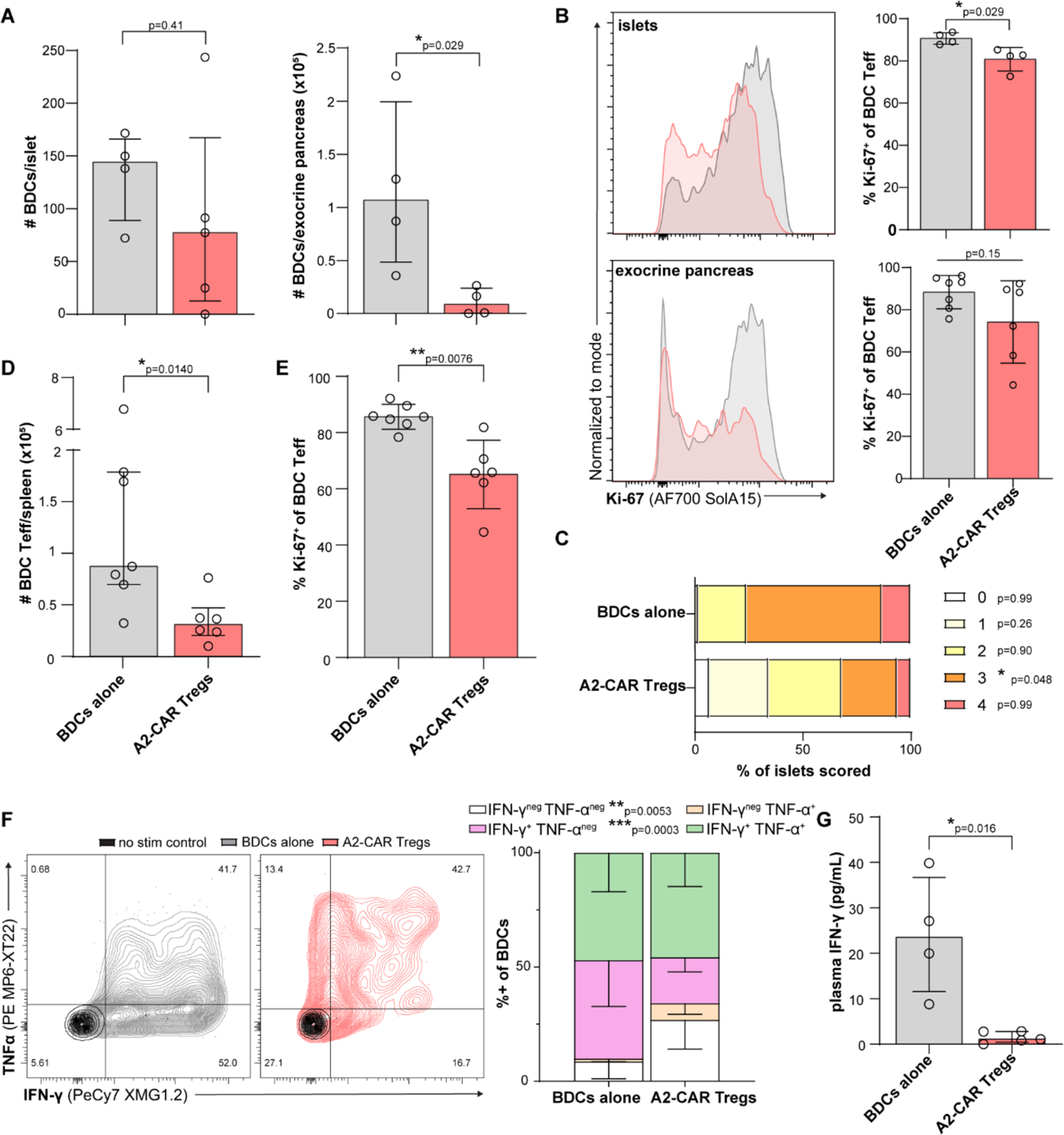
A2-CAR Tregs suppress BDC2.5 T cell effector function. **(A)** The number of BDC2.5 T cells found in the endogenous pancreas per islet (left) and in the exocrine pancreas (right) 7 days post-T cell injection (n=4-5; median with interquartile range; Mann-Whitney test). **(B)** Representative and averaged Ki-67 expression on BDC2.5 T cells in the endogenous islets (top) and exocrine pancreas (bottom) at day 7 (n=4-7; mean with SD; Welch’s t-test). **(C)** Pancreas sections taken on day 7 were assessed for insulitis (0 no infiltration – 4 high infiltration; n=2-3 mice per group; means graphed; 2-way ANOVA with Šídák’s multiple comparisons). **(D)** The number of BDC2.5 Teff per spleen 7 days post-T cell injection (n=6-7; median with interquartile range; Mann-Whitney test). **(E)** Ki-67 expression on BDC2.5 Teff in the spleens at day 7 (n=6-7; mean with SD; Welch’s t-test). **(F)** 7 days post-T cell injection, splenocytes were stimulated with the BDC2.5 mimetope peptide (P63) for 24 hours and intracellular IFN-γ and TNF-α were determined (n=9-11; mean ± SD; 2-way ANOVA with Šídák’s multiple comparisons). **(G)** IFN-γ in plasma collected on day 7 and pooled from multiple mice was measured. (n=13-14 mice; median with interquartile range; Mann-Whitney test).

To assess whether A2-CAR Treg treatment had a systemic impact on BDC2.5 T cell effector function, splenocytes were harvested on day 7 and stimulated with their mimetope P63 peptide. We found that A2-CAR Treg treated mice had a significant reduction in the proportion of BDC2.5 T cells producing IFN-γ (Fig 3F) and nearly undetectable levels of plasma IFN-γ, with a significant reduction compared to non-Treg treated mice (Fig 3G). Taken together, these data show that A2-CAR Treg treatment systemically suppresses BDC2.5 T cell engraftment, proliferation and inflammatory potential.

### A2-CAR Tregs suppress passenger HLA-A2^+^ T cells

A hallmark of autoimmune diabetes in NOD mice is islet infiltration of immune cells, especially autoimmune CD4^+^ and CD8^+^ T cells (*36*). We therefore considered the possibility that the transplanted islets from pre-diabetic, immunocompetent HLA-A2^+^ NOD mice may contain passenger T cells. Indeed, we found that all islet transplant recipients had detectable circulating HLA-A2^+^ CD4^+^ and CD8^+^ T cells prior to injection of BDC2.5 T cells, although their absolute numbers were highly variable (Fig 4A). Some transplanted mice were euthanized prior to adoptive T cell transfer due when passenger T cell engraftment alone (without addition of BDC2.5 T cells) was sufficient to induce hyperglycemia (data not shown). Remarkably, A2-CAR Treg treatment almost entirely eliminated these passenger T cells from the mouse. Seven days after T cell injection, HLA-A2^+^ T cells were almost undetectable in spleens (Fig 4B), and regardless of their pre-CAR-Treg treatment numbers in the blood, HLA-A2^+^ CD8^+^ and CD4^+^ T cells were significantly reduced in the circulation of A2-CAR Treg-treated, but not untreated or polyclonal Treg-treated, mice (Fig 4C). Similar to other tissues, there was a trend to fewer HLA-A2^+^ T cells in the endogenous islets of A2-CAR Treg-treated mice (Fig 4D). These data suggest that in this model, autoimmunity was likely caused by a combination of natural autoreactive T cells with multiple specificities and BDC2.5 T cells. Thus, A2-CAR Tregs mediated bystander and/or linked suppression to control multiple autoreactive specificities.

**Fig. 4:**
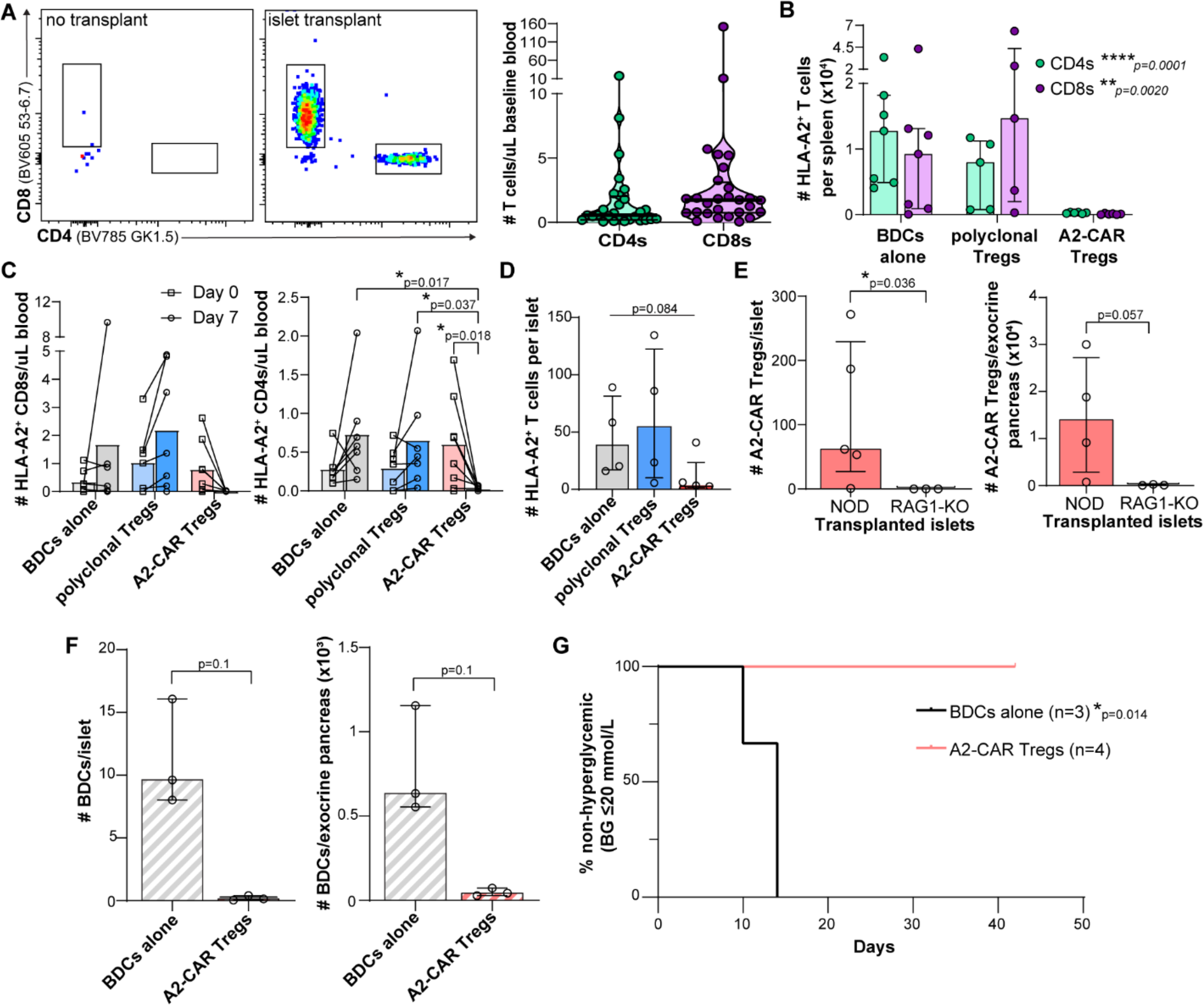
A2-CAR Tregs suppress islet-resident, HLA-A2^+^ passenger T cells. **(A)** Representative CD4^+^ and CD8^+^ T cells detected in blood in mice without or with an HLA-A2^+^ islet transplant. The adjacent violin plot shows the absolute number of HLA-A2^+^ T cells post-HLA-A2^+^ islet transplant prior to T cell injection (n=28; median with interquartile range). The number of HLA-A2^+^ CD4^+^ and CD8^+^ T cells in **(B)** spleens at day 7 (n=4-7; median with interquartile range; Kruskal-Wallis tests), and **(C)** blood on days 0 and 7 (n=7-8; individual pairs shown with bars at means; 2-way repeated measures ANOVA with Tukey’s multiple comparisons). **(D)** The number of HLA-A2^+^ T cells in the endogenous HLA-A2^negative^ islets 7 days post-T cell injection (n=4-7; median with interquartile range; Kruskal-Wallis tests). **(E-G)** Islets from HLA-A2^+^ NOD.Rag1-/-mice were transplanted into NSG mice, then injected with BDC2.5 T cells alone or with A2-CAR Tregs. **(E)** The number of A2-CAR Tregs found per endogenous islet and in the exocrine pancreas in mice transplanted with either HLA-A2^+^ NOD (NOD) or HLA-A2^+^ NOD.Rag1-/-(RAG1-KO) islets at day 7 (n=3-5; median with interquartile range; Mann-Whitney test). **(F)** BDC2.5 T cell accumulation in endogenous islets (left) and exocrine pancreas (right) at day 7 (n=3; median with interquartile range; Mann-Whitney test). **(G)** Diabetes-free survival of each treatment group transplanted with HLA-A2^+^ Rag1-KO islets (n=8-11; Mantel-Cox Logrank test).

The systemic presence of HLA-A2^+^ T cells raised the possibility that the effects of A2-CAR Treg treatment on BDC2.5 T cells were not mediated through linked suppression, but rather bystander suppression: e.g. potentially A2-CAR Tregs were stimulated by HLA-A2^+^ passenger T cells near BDC2.5 T cells, causing bystander suppression of the BDC2.5 T cells that was continuously dependent on A2-CAR Treg local presence and activation. To determine if the systemic tolerogenic effects of A2-CAR Tregs depended on the presence of passenger HLA-A2^+^ T cells, transplants were repeated using islets from A2^+^ NOD.Rag1-/-mice, which do not have any T or B cells. The absence of HLA-A2^+^ passenger T cells resulted in a significant reduction the number of A2-CAR Tregs in the endogenous pancreas (Fig 4E), but the accumulation of BDC2.5 T cells in the endogenous pancreas was nevertheless significantly reduced relative to untreated mice (Fig 4F). Moreover, A2-CAR Treg therapy completely protected these mice from diabetes (Fig 4G). These data show that the systemic protective effect of A2-CAR Tregs does not depend on HLA-A2^+^ T cell engraftment and presence of the CAR antigen in the endogenous islets. Thus A2-CAR Tregs may mediate linked suppression by modulating BDC2.5 T cell activity such that the suppressed effect no longer depends on the local presence of HLA-A2.

### A2-CAR Tregs induce infectious tolerance

To further explore whether A2-CAR Tregs had stably re-shaped BCD2.5 Teff activity, 4 weeks after treatment with A2-CAR Tregs we removed the cellular source of co-expressed A2-CAR and BDC2.5 T cell antigens by enucleating the eye previously transplanted with HLA-A2^+^ islets. As a control, some mice that received A2-CAR Tregs had their non-transplanted eye removed. We found that all enucleated mice maintained euglycemia for at least three weeks post-removal (Fig 5A). Removal of the HLA-A2^+^ graft did not result in a significant change in BDC2.5 Teff engraftment (Fig 5B). In the presence or absence of CAR antigen, the number (Fig 5C) and proportion (Fig 5D) of circulating FOXP3-expressing A2-CAR Tregs reached homeostatic levels and retained a level of de-methylation at the Treg-specific demethylation region (TSDR) consistent with stable Treg lineage (Fig 5E).

**Fig. 5:**
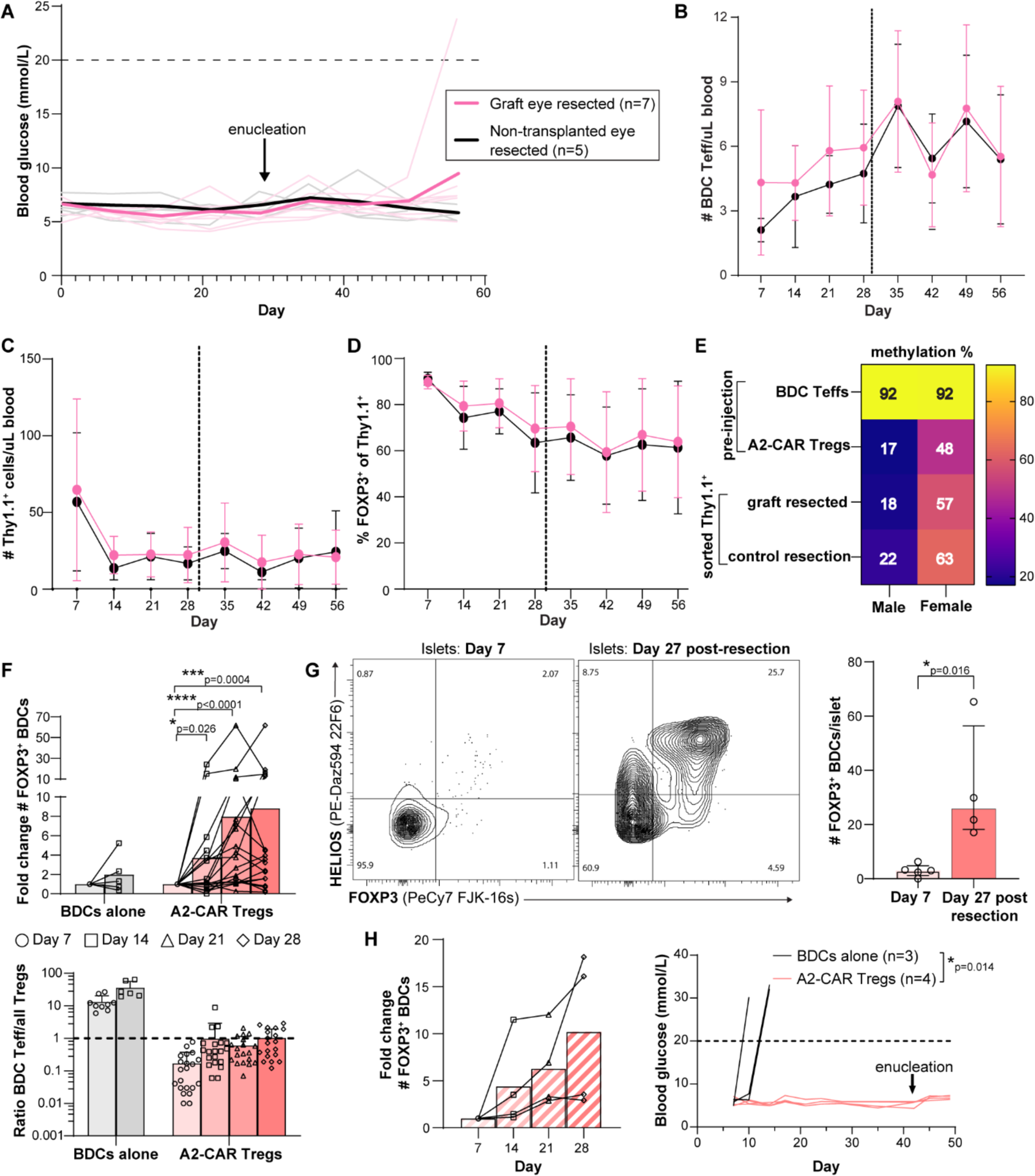
A2-CAR Tregs induce stable islet tolerance and FOXP3^+^ BDC2.5 T cells. **(A)** Twenty-eight days post-A2-CAR Treg injection, mice had either their non-transplanted (black) or islet-transplanted eye (pink) removed and blood glucose was monitored for another 28 days. Light lines show individual mice and the heavy lines show mean blood glucose readings for each group. **(B)** BDC2.5 Teff and Thy1.1^+^ A2-CAR Treg **(C)** engraftment and **(D)** FOXP3 expression in the blood over time leading up to and following enucleation, indicated by vertical dashed line (n=5-7; median +/-SD). **(E)** Four to six weeks following enucleation, splenic Thy1.1^+^ cells were sorted for TSDR analysis. BDC2.5 Teff and A2-CAR Treg methylation prior to cell injection are shown in the top two rows as controls (mean of n=1-4). **(F)** Top graph: fold change in numbers of FOXP3^+^ BDC2.5 T cells in blood at the indicated times. For mice with 0 Tregs on day 7, day 14 was considered the baseline (n=6-21; bars at mean fold-changes; Wilcoxon signed rank test where theoretical median = 1). Bottom graph: ratio of FOXP3^negative^ BDC2.5 Teff to the sum of FOXP3^+^ cell (BDC + Thy1.1) (n=6-21; mean ± SD). **(G)** Representative flow plots and averaged data of FOXP3 expression of islet-infiltrating BDC2.5 T cells in A2-CAR Treg-treated mice 7 days post-T cell injection or 27 days post-enucleation (n=4; median with interquartile range; Mann-Whitney test). **(H)** In mice transplanted with HLA-A2^+^ NOD.Rag1-/-islets and treated with A2-CAR-Tregs: left, the fold change of FOXP3^+^ BDC2.5 T cells in blood (n=4; paired samples with bars at mean fold-changes) and right, blood glucose levels before and after removal of the transplanted eye at day 42 (n=3-4; Mantel-Cox Logrank test).

Building upon evidence that A2-CAR Treg treatment significantly diminished BDC2.5 T cell effector function, we next sought evidence for infectious tolerance. We first assessed the fold change in the number of circulating FOXP3^+^ BDC2.5 T cells over time. Whereas untreated mice had no significant increase in circulating FOXP3^+^ BDC2.5 T cells from day 7 to 14, in A2-CAR Treg treated mice they significantly increased each week following day 7 (Fig 5F, top). Ultimately, this increase in FOXP3^+^ BDC2.5 T cells contributed to the emergence of a long-lasting, tolerogenic balance of BDC2.5 Teff to total Tregs (A2-CAR plus BDC2.5 Tregs) in A2 CAR Treg-treated mice that was never achieved in untreated mice (Fig 5F, bottom).

Moreover, there was evidence that FOXP3^+^ BDC2.5 T cells expanded in endogenous islets over time. Seven days following A2-CAR Treg injection, the proportion of FOXP3^+^ BDC2.5 T cells in endogenous islets was low (mean FOXP3^+^ =5.4%, Fig 5G), but overtime this population expanded ∼five-fold such that 27 days post-enucleation, about one third of BDC2.5 T cells resident in islets expressed FOXP3 (mean FOXP3^+^ = 27.7%, Fig 5G), indicating infectious tolerance. Importantly, we found a similar trend in A2-CAR Treg-treated mice transplanted with A2^+^ NOD.Rag1-/-islets; these mice also had an increase in circulating BDC2.5 Tregs (Fig 5H, left) and maintained euglycemia post HLA-A2^+^ graft removal (Fig 5H, right). Thus, even in the absence of the A2-CAR antigen, BDC2.5 T cells were rendered tolerant to the endogenous pancreas by A2-CAR Treg treatment.

## DISCUSSION

Taken together, our data suggest that A2-CAR Tregs suppress islet-autoreactive T cells, mediating both linked suppression and infectious tolerance. These findings have significant implications for the field of Treg therapy because they provide evidence that an engineered Treg product has the potential to permanently re-shape immune responses such that long-term tolerance is maintained by cells other than the therapeutic Tregs.

To the best of our knowledge, these are the first data to formally show that an antigen-engineered Tregs can induce long-lasting tolerance to a distinct antigen *in vivo* via linked suppression. Although widely cited as a key Treg mechanism of action, there have been few *in vivo* studies showing evidence of linked suppression. There is the original observation that tolerance to an “A” skin graft results in tolerance to co-expressed “B” antigens on another skin graft (“AxB” skin) even in the subsequent absence of “A” (*18*). There is also the finding that autoantigen-detecting Tregs isolated from TCR-trangenic mice reduced symptom severity in a mouse model of multiple sclerosis elicited by a distinct autoantigen (*37*). In terms of bystander suppression, multiple reports show engineered Tregs suppress diabetogenic T cells with distinct antigen specificities *in vitro* (*28, 38, 39*). *In vivo*, MHC:insulin-specific CAR Tregs limited BDC2.5 T cell engraftment (*28*) and MHC class I-specific CAR Tregs protected multiple transplanted tissues from alloreactive immunity (*40*). Although these studies demonstrated bystander suppression, they did not show long-lived tolerance to distinct antigens even in the absence of the CAR antigen. Our model uniquely allowed us to evaluate the capacity of A2-CAR Tregs to inhibit BDC2.5 effector function both in the presence (bystander) and absence (linked) of A2-CAR antigen expression. We found that in the endogenous HLA-A2^negative^ pancreas, in the absence or presence of HLA-A2^+^ passenger T cells, BDC2.5 T cells had suppressed engraftment, proliferation and inflammatory potential even if there were no A2-CAR Tregs in the vicinity. Moreover, A2-CAR Treg treatment not only suppressed BDC2.5 effector function, it resulted in a re-balanced, tolerant immune state.

We found that the endogenous HLA-A2^negative^ pancreas was protected and functional even after HLA-A2^+^ graft removal, an outcome correlated with a significant increase in FOXP3^+^ BDC2.5 T cells residing in endogenous islets. These data suggest that A2-CAR Tregs tolerized BDC2.5 T cells in a long-lasting way, demonstrating infectious tolerance. This finding has significant implications for the engineered Treg therapy field because if these cells can permanently induce tolerance, their continued presence may not be required. A limitation is that we were unable to completely eliminate all FOXP3^+^ BDC2.5 T cells prior to injection so we cannot determine whether the increased population of FOXP3^+^ BDC2.5 Tregs arose from preferential expansion of a small number of transferred BDC2.5 Tregs and/or via *de novo* conversion into new Tregs. Nevertheless, in either case the ability of A2-CAR Tregs to expand FOXP3^+^ BDC2.5 T cells is a significant advancement in our understanding of how Treg therapy can re-shape tolerance.

A2-CAR Tregs reduced engraftment of diabetogenic BDC2.5 T cells and islet-resident passenger T cells. Mechanistically, Tregs are thought to mediate suppression via APC modulation, tolerogenic cytokine release and metabolic microenvironment manipulation. There is also some evidence that Tregs may engage in cytolysis via granzyme/perforin and/or Fas/FasL pathways (*41*). Our findings raise the question of where A2-CAR Tregs mediated their effects. Although A2-CAR Tregs suppress DC function (*16*), NSG mice have dysfunctional DCs and small, hypocellular lymphoid organs (*42*), so in this model, A2-CAR Tregs most likely suppress BDC2.5 T cells at the site of transplantation, resulting in a systemic effect. This possibility is supported by intravital imaging studies of mice with ACE islet transplants and injected with BDC Teff and Tregs which showed that Tregs create long-lasting bonds with BDC2.5 T cells, more so than with DCs, in a partially CTLA-4-dependent manner (*33*).

Although the BDC2.5 transfer model of autoimmune diabetes has caveats, it uniquely allowed us to probe the questions of bystander/linked suppression and infectious tolerance. The BDC transfer model uniquely allowed us to evaluate the impact Treg therapy on autoimmune CD4^+^ T cells in the absence of a confounding anti-HLA-A2 alloimmune response which may be difficult to parse from the autoimmune response. Another advantage of the BDC transfer model was that the lymphodeplete NSG recipients already had “space” for Treg engraftment, to a degree mimicking the situation in people undergoing islet transplantation receiving aggressive conditioning regimens in preparation (*26*). T1D is an autoimmune disorder in which CD4^+^ T cells, CD8^+^ T cells and B cells all have pathogenic roles (*43*), and our inability to evaluate the impacts of CAR Tregs on these cells is a limitation of this study. A future direction is to test A2-CAR Tregs in an immunocompetent NOD model of T1D. However, given the small number and polyclonal repertoire of autoimmune T cells in NOD mice, precise tracking the impact of Tregs on these naturally developed cells is challenging.

Overall, the results of this study show that antigen-specific CAR Tregs can be activated *in vivo* to induce tolerance of autoantigens via linked suppression. The first in-human trials of A2-CAR Tregs in transplantation are underway (NCT04817774, NCT05234190) and will provide important safety and efficacy data. Upon further validation of A2-CAR Treg efficacy and infectious tolerance induction in immunocompetent preclinical models of autoimmunity, a natural next step for A2-CAR Treg therapy in humans will be use in transplantation with the intent to control both allo and autoimmunity.

## MATERIALS AND METHODS

### Study design

We developed an *in vivo* model of islet transplantation with autoimmune diabetes challenge to test whether CAR Tregs that detect an alloantigen were able to suppress autoantigen-specific CD4^+^ T cell-induced T1D when both target antigens were expressed by the same tissue (the islet transplant). All data were generated using mouse immune cells. After figure 1, which featured Treg phenotyping and an *in vitro* co-culture experiment, all data were derived from *in vivo* mouse experiments. NSG mice were transplanted with HLA-A2^+^ islets and then administered BDC2.5 Teff to induce T1D. Mice received BDC2.5 Teff alone or were co-administered with polyclonal or A2-CAR Tregs (3:1 BDC2.5 Teff). Experimental treatments were distributed evenly among transplanted mice, including sexes and cage mates. Investigators were not blinded to experimental groups, with the exception of insulitis rating. Cohort sizes (approximately 12 mice) were restricted by islet donors and yield for transplant. Most experiments were repeated with at least two transplanted cohorts, with the exception that Rag1-KO transplant experiments were performed once with an n of 3-4 per experimental condition. Blood glucose was regularly monitored for hyperglycemia (≥20 mM glucose). Immune populations were monitored weekly in the blood and in the spleen and pancreas at endpoint. Mice were euthanized once hyperglycemic, 7 days post-T cell injection or 4-6 weeks post-last intervention.

### Animals and ethics

All *in vivo* experiments were approved by the University of British Columbia Animal Care Committee (A20-0017). NSG (JAX 005557) and NOD.FOXP3-GFP (JAX 008694) were purchased from The Jackson laboratory and bred in-house under specific pathogen-free conditions. NOD-HLA-A2 mice were generated by crossing the HLA-A2 from NSG-HLA-A2 (JAX 014570) onto the NOD/ShiLtJ (JAX 001976) background, resulting in mice with wildtype Prkdc, wildtype IL2rg, and homozygous HLA-A2. To generate NOD.Rag1KO-HLA-A2 mice, NOD.Rag1KO-MIP.CFP mice received from Dr. Brain Fife (University of Minnesota) were crossed with NOD-HLA-A2 mice until progeny were homozygous for Rag1-KO, and transgenic positive for MIP-CFP (cyan fluorescent protein expressed under the mouse insulin 1 promoter) and HLA-A2. NOD.FOXP3-GFP x BDC2.5 mice were generated by crossing NOD.BDC2.5 (JAX 004460) and NOD.FOXP3-GFP mice and subsequently genotyped for FOXP3-GFP and BDC2.5 TCR co-expression.

### Flow cytometry

Dead cells were identified and excluded based on extracellular staining with fixable viability dye (FVD eFluor 780, eBioscience). Cells that received an intracellular stain were fixed and permeabilized using the FOXP3/Transcription Factor staining buffer set (Thermo Fisher Scientific). Flow cytometry data were collected on the LSRFortessa X-20, A5 Symphony (BD Biosciences) or Cytoflex (Beckman Coulter). Data were analyzed with FlowJo version 10.

### Treg generation

Treg generation from NOD.FOXP3-GFP mice was previously described (*28*). Splenocytes and lymph nodes were collected from adult (age ≥ 8 weeks) female or male NOD.FOXP3-GFP mice. Lymphoid organs were homogenized in 15 mL dounce tissue grinders (VWR) and CD4+ T cells were enriched via magnetic negative selection (STEMCELL Technologies). Tregs were then sorted on a BD FACS Aria Fusion based on viability dye^negative^ CD4^+^ (BV605, clone RM4-5, BD) GFP-FOXP3^+^. Tregs were seeded at an approximate density of 1.2 million cells/cm^2^ in ImmunoCult-XF T cell expansion medium (STEMCELL Technologies) supplemented with 50 μM β-mercaptoethanol (Sigma-Aldrich) and penicillin-streptomycin (Gibco), with 1000 U/mL IL-2 (Proleukin), 50 nM rapamycin (Sigma-Aldrich) and a 3:1 CD3/CD28 mouse T-activator Dynabead to Treg ratio (Thermo Fisher Scientific). On day 3 of expansion, Tregs were transduced with retrovirus, 2 μg/ml Lipofectamine 2000 (Thermo Fisher Scientific) and 1.6 μg/mL Polybrene (MilliporeSigma) by centrifuging at 32°C and 805 xg for 90 minutes. On day 7, Dynabeads were magnetically removed and Thy1.1+ cells were positively selected (STEMCELL Technologies). Tregs rested overnight until day 8, at which point they were phenotyped (FOXP3-PeCy7, clone FJK-16s, Thermo Fisher Scientific; HELIOS-PEDazzle594, clone 22F6, Biolegend; Thy1.1-BV711, clone OX-7, BD Biosceinces; myc-Alexa Fluor 647, clone 9E10, UBC abLab) and frozen for future use.

### DC suppression assay

This assay has been previously described (*16*). Spleens were collected from adult, sex-matched NOD/ShiLtJ or NOD-HLA-A2 mice and enzymatically digested (STEMCELL Technologies). CD11c^+^ DCs were positively magnetically isolated (STEMCELL Technologies). DCs were plated at a 1:2 ratio with Tregs and left to co-incubate for 2 days, at which point CD11c^+^ (SB780, clone N418, BD Biosciences) DC expression of CD80 (BUV395, clone 16-10A1, BD Biosciences), CD86 (BV650, clone GL1, BD Biosciences) and I-A^g7^ (PacBlue or AF488, clone 10.2-16, provided by Dr. Brian Fife at the University of Minnesota) were measured by flow cytometry.

### Islet isolation and transplantation

Islets were isolated from male or female NOD-HLA-A2 (age 6-11 weeks) or NOD-Rag1KO-HLA-A2 mice (age ≥ 8 weeks). The pancreas was exposed in euthanized mice and the common bile duct’s entry to the duodenum clamped. The common bile duct was cannulated with a 27G needle and the pancreas inflated with 1000 U/mL of collagenase XI (Sigma-Alrich) in Hank’s Balanced Salt Solution (Gibco). The pancreas was excised and incubated in a 39°C water bath for 15 minutes. The pancreas was gently shaken to dissociate islets from the exocrine pancreas. The islets were washed and caught on a 100 μm filter, and then rinsed into a non-tissue culture treated petri dish in RPMI (Gibco), supplemented with 10% fetal bovine serum (Gibco, Seradigm, VWR), penicillin-streptomycin and glutamax (Gibco). Islets were hand-picked and transplanted on the same or following day.

NSG mice (age 28 weeks) were transplanted with 50-150 islets as previously described (*30*). Given that the NSG mice were incapable of raising an allogeneic immune response targeting the islets, sex-matching was preferred but not required. We did not observe any impact of sex on study results. Once under anesthesia, a 27G needle was used to create a small hole entering into the ACE of the left eye. This hole was slightly enlarged using a pair of fine scissors and the islets, collected in a stripper (Origio MidAtlantic Devices) connected to a micromanipulator, were carefully placed into the ACE.

### Longitudinal metabolic and immunological monitoring

Blood glucose was measured via saphenous bleed using the OneTouch Ultra 2 machine and glucose strips. Blood glucose was measured starting seven days post-BDC2.5 Teff injection, and then 1-2 times a week until endpoint. Mice were deemed diabetic when their blood glucose measured 20 mM or above. Blood was collected from the saphenous vein for flow analysis once a week. Blood was spun for 10 minutes at 450 xg and plasma volume measured and frozen at - 20°C for later analysis. The remaining cellular pellet volume was measured and the red blood cells lysed with ammonium chloride (STEMCELL Technologies). Cells were then stained to identify viability dye^negative^ CD45^+^ (BUV395, clone 30-F11, BD Biosciences) leukocytes. Adoptively transferred Tregs were further identified as Thy1.2^+^ (BV510, clone 53-2.1, Biolegend) CD4^+^ (BV785 or BV605, GK1.5 or RM4-5, BD) Thy1.1^+^. BDC2.5 Teff were identified as Thy1.2^+^ Thy1.1^negative^ CD4^+^ Vβ4^+^ (FITC, clone KT4, BD Biosciences) P63 tetramer^+^ (PE, provided by Dr. Brian Fife’s laboratory, University of Minnesota). Passenger T cells were identified as CD4^+^ or CD8^+^ (BUV805 or BV605, clone 53-6.7, BD Biosciences) and HLA-A2^+^ (PE, APC or BV421, clone BB7.2, Biolegend, BD Biosciences or Invitrogen). Cells were stained intracellularly for FOXP3, HELIOS and Ki-67 (AlexaFluor 700 or eF450, clone SolA15, Invitrogen). Total cell counts were calculated based on total blood volume collected and 123count eBead flow cytometry controls (Thermo Fisher Scientific).

### Diabetes induction and Treg treatment

Spleens and lymph nodes were collected from male and female NOD.FOXP3-GFP x BDC2.5 mice, homogenized to single cell suspensions and magnetically enriched for CD4^+^ T cells as already described. BDC Teff were then sorted as viability dye^negative^ CD4^+^ FOXP3-GFP^negative^. Cells were immediately frozen for later use. At least two weeks following islet transplantation, the blood glucose of transplant recipients was measured at baseline to exclude hyperglycemic (≥20 mM) mice (occasionally induced by passenger T cells). BDC2.5 Teff and Tregs were thawed, counted and injected via tail vein in a volume of 150μL PBS. Mice transplanted with NOD-HLA-A2 islets were injected with 50,000 BDC Teff ± 150,000 Tregs in the same injection. BDC-induced diabetes was less effective in the absence of passenger T cells, so mice transplanted with NOD-Rag1KO-HLA-A2 islets were injected with 150,000 BDC Teff ± 450,000 Tregs, maintaining the same Teff:Treg ratio.

### Plasma cytokine analysis

Plasma samples isolated from blood collected at day 7 and previously frozen at -20°C were thawed and pooled to reach the minimum sample volume required for cytokine analysis (20μL) using the Th1/Th2/Th17 cytokine kit (BD Biosciences). Data were analyzed using the FCAP array software, version 3.0.1 (Soft Flow).

### Endpoint phenotyping

On day 7, splenocytes were harvested as already described. Islets were collected as described, and flow through from islet isolation was kept and analyzed as exocrine pancreas. Absolute number of cells are reported for whole spleen and all exocrine tissue analyzed. Islets were counted and hand picked before gently trypsinizing at 37°C to achieve single cell suspensions. Absolute counts for islets are reported per islet (total number of cells/number of hand-picked islets).

### Insulitis scoring

On day 7, pancreases were harvested, gently fixed at 4°C for 2-4 hours in Cytofix (diluted 1:4 in PBS; BD biosciences), then embedded in O.C.T. embedding compound (Fisher Scientific or Sakura) and frozen at -80°C. Four 12 μm sections spaced 50 μm apart were fixed in 95% Ethanol and then stained with Hematoxylin and Eosin. A blinded insulitis score was applied to each islet; ‘0’ represented no immune cells surrounding the islet, ‘1’ represented peri-insulitis (mononuclear cells surrounding the islet periphery along with periductal insulitis), ‘2’ represented moderate insulitis (mononuclear cells infiltrating <50% of islet area), ‘3’ represented severe insulitis (mononuclear cells infiltrating >50% of islet area), and ’4’ represented severe structural derangement of the islet (mononuclear cells infiltrating center of the islet with few remaining cells).

### Splenocyte intracellular cytokine staining

At day 7, spleens were harvested and homogenized into single cell suspensions. Red blood cells were lysed using ammonium chloride solution (STEMCELL Technologies). Five hundred thousand splenocytes were plated with 500 nM of an irrelevant peptide (hen egg lysozyme HEL_11-25_: AMKRHGLDNYRGYSL) or the BDC2.5 mimetic peptide (P63: RTRPLWVRME) (GenScript). After 20 hours of co-incubation, 10 μg/mL of Brefledin A (Sigma-Aldrich) was gently added. After a total of 24 hours, cells were stained intracellularly for IFN-γ (PE-Cy7, clone XMG1.2, eBiosciences) and TNF-α (PE, clone MP6-XT22, BD Pharmingen).

### Enucleation

To test whether the endogenous pancreas was functional four weeks post BDC2.5 and A2-CAR Treg injection, either the islet graft-containing eye (left) or the non-transplanted control eye (right) was enucleated under gaseous isoflurane. Once in the surgical level of anesthesia, the eye was gently lifted out of the eye socket and two sutures were tied around the optical nerve. The eye was then cut above the suture ties and the eyelids glued shut with Vetbond tissue adhesive (3M).

### Treg specific demethylated region (TSDR) analysis

Four to six weeks post-enucleation, spleens were harvested and homogenized into single cell suspensions. Thy1.1^+^ Tregs were sorted on the BD FACS Aria Fusion based on viability dye^negative^ CD4^+^ Thy1.1^+^. Spare BDC2.5 Teff and A2-CAR Tregs thawed for T cell injection were used as controls. Genomic DNA was isolated from previously frozen cell pellets and bisulfite converted using the EZ DNA methylation-direct kit (Zymo Research). Two PCRs were performed with the PyroMark PCR kit (Qiagen) in order to report the average methylation of 8 representative CpG sites in the regulatory T cell–specific demethylated region (TSDR) on the X chromosome. Pyrosequencing was performed on a PyroMark Q96 ID (Qiagen) with PyroMark Gold Q96 reagents (Qiagen) and Streptavidin Sepharose (GE Healthcare). Results for male and female T cells are represented separately and unadjusted; i.e. reported female methylation has not been adjusted to account for X inactivation. PCR primers: PCR1 FWD – TGGGTTTTGTATGGTAGTTAGATGG, PCR1 REV-Biotin – CCCTATTATCACAACCTAAACTTAACC, SEQ1 FWD – TTGTATGGTAGTTAGATGGA, PCR2 FWD-Biotin – GGGTTTTGTATGGTAGTTAGATGG, PCR2 REV – AAACCCTATTATCACAACCTAAACTT, and SEQ2 REV – CCTAAACTTAACCAAATTTTTCT.

### Statistical analyses and figure preparation

Figure schematics were generated in BioRender. Graphs were generated with GraphPad Prism 10. Statistical testing was also completed in GraphPad Prism 10; the statistical test used for each panel is described in the figure legend. Generally, reports of cell numbers are presented as medians with interquartile range and assumed non-parametric due to wide variance in engraftment, while frequencies of marker expression are presented as means with standard deviations (SD) and assumed parametric. Plots that show paired values have bars at the means.

### List of Supplementary Materials

*None*.

## Acknowledgments

We thank Dr. Lisa Xu for her aide with fluorescence-activated cell sorting, the animal care staff a the BC Children’s Hospital Research Institute and Centre for Molecular Medicine and Therapeutics, and Melissa Braschel for statistical guidance.

## Funding

Helmsley Charitable Trust 2018PG-T1D058 and G-2303-06775 (MKL, BTF)

Canadian Institutes for Health Research PJT-495696 (MKL)

National Institutes of Health R01 AI156276 (BTF)

Canadian Graduate Scholarship – Doctoral (CMW)

BC Children’s Hospital Research Institute (MKL)

Canada Research Chair in Engineered Immune Tolerance (MKL)

## Author contributions

Conceptualization: CW, MKL

Methodology: CW, VF, MM, MKL

Investigation: CW, VF, EC, MH, JG, MM

Visualization: CW

Funding acquisition: BTF, MKL

Project administration: CW, MKL

Supervision: MKL

Writing – original draft: CW, MKL

Writing – review & editing: CW, VF, JAS, MM, BTF, MKL

## Competing interests

MKL holds patents and a license related to A2-CAR technology. All other authors declare that they have no competing interests.

## Data and materials availability

All data are available in the main text.

